# Enhancer redundancy predicts gene pathogenicity and informs complex disease gene discovery

**DOI:** 10.1101/459123

**Authors:** Xinchen Wang, David B. Goldstein

## Abstract

Non-coding transcriptional regulatory elements are critical for controlling the spatiotemporal expression of genes. Here, we demonstrate that the number of bases in enhancers linked to a gene reflects its disease pathogenicity. Moreover, genes with redundant enhancer domains are depleted of *cis*-acting genetic variants that disrupt gene expression, and are buffered against the effects of disruptive non-coding mutations. Our results demonstrate that dosage-sensitive genes have evolved robustness to the disruptive effects of genetic variation by expanding their regulatory domains. This resolves a puzzle in the genetic literature about why disease genes are depleted of *cis*-eQTLs, suggesting that eQTL information may implicate the wrong genes at genome-wide association study loci, and establishes a framework for identifying non-coding regulatory variation with phenotypic consequences.

## Introduction

Non-coding regulatory elements, such as transcriptional enhancers, are critical for the precise spatiotemporal regulation of gene expression. Transcriptional regulation is a highly complex process often mediated by arrays of enhancer elements separated from their regulated genes by over one megabase^1^. Studies in *Drosophila* have demonstrated that key developmental genes are often regulated by multiple “shadow” enhancers with redundant activity patterns to protect against genetic perturbations^1–3^. Recent work suggests that a similar organizational structure may be present in mammalian genomes. Mammalian developmental genes have been reported to be near more enhancer elements than the average gene^4,5^, and targeted deletions of ultraconserved enhancers regulating key developmental genes has led to viable mice, occasionally with subtle abnormal phenotypes^6–8^. Moreover, differences in activity of enhancer elements are often not reflected by changes in gene expression^9–11^.

In keeping with the key role that non-coding enhancer elements play in gene regulation, many groups have recently shown that enhancers are enriched for disease-associated common genetic variants^12–14^. In contrast, disease-associated rare variants have mostly been identified within the protein-coding regions of genes, and studies employing whole-exome sequencing (WES) have effectively implicated genes and disease-causing mutations in conditions including epilepsy, idiopathic pulmonary fibrosis, ALS, and others^15–17^. Despite intense interest in developing complementary approaches to implicate rare non-coding disease mutations, progress has been limited, and many studies employing whole-genome sequencing (WGS) have failed to discover equivalently large sets of rare variant signals in non-coding regions^18–21^. Our limited understanding of enhancer biology is a major roadblock in the successful application of WGS for disease diagnosis. In particular, disruptive non-coding mutations are difficult to recognize, as the specific functional nucleotides within enhancers remain unknown^22–24^. The target genes of enhancers are also poorly resolved, as experimental methods have limited resolution and/or limited throughput, while computational methods often suffer from poorly understood accuracy^23,25–28^. These limitations complicate the systematic study of enhancers in rare disease genetic mapping studies. Thus, the development of a framework to study enhancers in human disease will have major implications for genetic mapping studies.

Inspired in part by the shadow enhancer model, we hypothesized that the disease pathogenicity of a gene could be predicted by its regulatory landscape. Here, we use computational predictions of enhancer-gene interactions to develop a simple scoring system to rank genes by the number of functional regulatory nucleotides in their enhancer elements. Remarkably, this scoring system, which we term the “enhancer domain score”, is highly reflective of gene pathogenicity and is independent of and complementary to existing metrics of intolerance and constraint. This result also provides a genome-wide assessment of the performance of computational methods for predicting which important enhancers regulate human genes. We show that the enhancer domain score negatively correlates with whether the associated gene carries a *cis*-acting eQTL, suggesting that mammalian disease genes have evolved a robustness to regulatory genetic variation, similar to past observations in *Drosophila*. Notably, candidate causal genes at genome-wide association study (GWAS) loci have high enhancer domain scores, suggesting these genes may be less likely to be discovered using eQTLs. Indeed, we provide evidence showing that eQTL information tends to implicate the wrong causal genes at these GWAS loci. Finally, we show that the identification of these enhancer regions of genes provides an appropriate framework for the identification of disease-causing mutations in regulatory sequences and emphasizes the importance of using approaches that assess the cumulative burden of genetic variation falling in implicated enhancer regions.

## Results

### Genes with large functional regulatory domains are associated with developmental Diseases

We hypothesized that the transcriptional regulatory landscape of a gene reflects properties of the gene itself. We therefore sought to construct an enhancer regulatory score for human genes and assess its ability to prioritize genes important in human disease (Fig. 1A). The association of *cis*-acting regulatory elements with their target genes remains an outstanding problem in the field of genomics and gene regulation^23,25^. While many approaches have been developed to address this challenge, most methods are currently constrained to a limited number of enhancer elements or tissue types. To generate a genome-wide compendium of transcriptional enhancer elements and their target genes, we initially relied on predictions based on correlation between predicted enhancer activity and gene expression from 127 human tissues profiled from the Epigenome Roadmap project using the approach developed by Ernst *et al*. and Liu *et al*. (“activity-linking”, Fig. 1B)^4,29^.

Under the shadow enhancer model, developmental genes in *Drosophila* have multiple redundant enhancers to buffer against deleterious regulatory mutations^2,3^. We therefore hypothesized that human genes with more linked enhancers are more important in mammalian development and, as a consequence, human disease. To test this hypothesis, we ranked genes by five metrics that reflect the size of their transcriptional regulatory elements:

i. The number of discrete enhancer elements linked to a gene
ii. The total number of nucleotides within all linked enhancers
iii. The total number of nucleotides that show evolutionary conservation within all linked enhancers (union of conserved elements across vertebrates, placental mammals and primates, see **Fig. S1**)
iv. The total number of nucleotides in cis-regulatory modules predicted by the UniBind database within linked enhancers^30^
v. The total number of nucleotides in transcription factor binding sites annotated by the UniBind database within linked enhancers^30^

Strikingly, genes highly ranked by these metrics are significantly enriched for haploinsufficient genes (**Fig. S2**), developmentally important genes in mouse (Fig. 1C), and genes deposited in the Online Mendelian Inheritance in Man (OMIM) database and linked to human disease (**Fig. S2**), suggesting a general biological principle whereby developmentally-important genes have larger functional regulatory domains. Among the five metrics tested, the total number of conserved nucleotides within all linked enhancers (metric #3) was consistently the most enriched metric for disease-relevant genes (Fig. 1C, **Fig. S2**). In subsequent analyses we use this metric as a proxy for the total number of functional elements in linked enhancers. We refer to this as a gene’s “enhancer domain score”, or “EDS” (list of genes by EDS available in **Table S1**, distribution of EDS for all genes shown in Fig. 1D).

We observed that genes ranking highly by EDS are enriched for many different disease gene sets, including OMIM, Developmental Disorders Genotype-Phenotype Database (DDG2P) genes^31^, and genes with a “likely pathogenic” or “confirmed pathogenic” ClinVar variant (Fig. 1E). These “high EDS genes” (top 3000 genes by EDS, cut-off chosen based on comparable numbers of human constrained genes, Methods) are also substantially more likely to be haploinsufficient and lead to embryonic lethality when knocked out in mouse. Furthermore, this enrichment persists over a range of EDS cut-offs and is strongest for genes with the greatest EDS (Fig. 1F,G). As a negative control, we note that high EDS genes are strongly depleted for olfactory receptors (Fig. 1E), consistent with previous reports that mutations in olfactory receptors are tolerated in humans and do not lead to developmental disease^31,32^. We also split DDG2P developmental disease genes by the affected organ, and observed that high EDS genes are enriched for developmental diseases affecting a wide range of human organs (Fig. 1H)^31^. This indicates that the EDS score is not driven by the effect of any individual tissue but that the relationship between the functional enhancer nucleotide count and gene pathogenicity may represent a fundamental biological principle.

**Figure 1.**
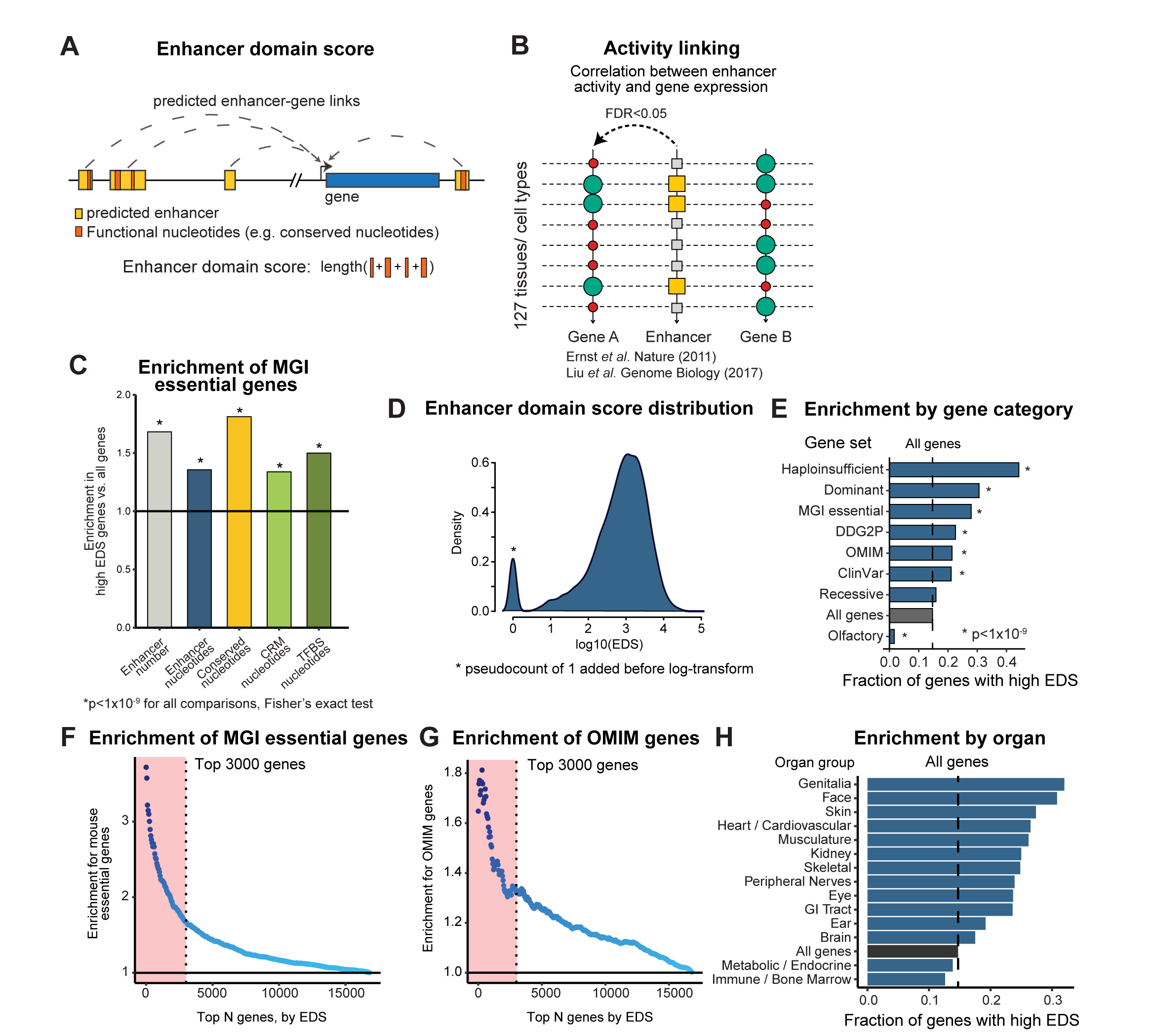
Enhancer domain score is associated with gene pathogenicity. **(A).** Overview of the approach to quantify the enhancer domain size for a gene based on linked enhancer elements. **(B)** Overview computational approach to linking enhancers to nearby genes (“Activity-linking”). Enhancer-gene links are assigned by the correlation between enhancer activity and gene expression across 127 human tissues. **(C)** Comparison of enrichment for MGI mouse essential genes using genes ranked highly by five regulatory metrics: total number of enhancers, total count of enhancer nucleotides, total count of conserved enhancer nucleotides, total number of nucleotides within predicted *cis*-regulatory modules (CRM), and total number of nucleotides within transcription factor binding sites (TFBS). The top 3000 genes ranked by each metric are used for the enrichment foreground set. Conserved enhancer nucleotides represents union of conserved nucleotides across vertebrates, placental mammals and primates **(D)** Distribution of enhancer domain scores using activity-linking to assign enhancers to nearby genes. **(E)** Genes ranking highly by the enhancer domain score are enriched for disease-relevant gene sets. Top 3000 genes used as foreground. **(F,G)** High EDS genes are enriched for MGI essential genes (panel F) and OMIM disease genes (panel G) across a wide range of cut-offs. Red area corresponds to top 3000 genes, used in all subsequent analyses as “high EDS genes”. **(H)** Enrichment of high EDS genes within developmental disease gene sets is high across disease genes that affect a variety of human organs.

To control for possible artifacts from the enhancer-gene linking approach used, we repeated the analyses presented above using an independent enhancer-gene linking approach (“proximity-linking”) that assigns enhancers to their nearest gene using the GREAT algorithm and does not consider enhancer activity patterns or gene expression^33^. We observed that genes ranking highly under the proximity-linking EDS are likewise enriched for the disease gene sets (**Fig. S3**), indicating the enrichment observed for activity-linking EDS is not due to artifacts from the activity correlation approach used above.

### The enhancer domain score provides information independent of population-based metrics of gene intolerance and constraint

In recent years, constraint-based metrics of gene essentiality have been developed that reflect the absence of loss-of-function (LoF) variants (pLI^32^) or the discordance between LoF, missense, and synonymous variants in the human population (RVIS^34^). Genes that rank highly by the pLI or RVIS metrics have been shown to be significantly more likely to be involved in human disease. These rankings are therefore commonly applied to prioritize causal genes from clinical genetics studies^35^. To investigate whether the enhancer domain score adds value beyond constraint-based gene metrics, we selected the top 3000 genes ranked by each method (EDS, pLI and RVIS), corresponding to a pLI cut-off of 0.918 and RVIS cut-off of 18.5%, and compared each set to identify similarities and differences in disease relevance. We used this cut-off so that each set has the same number of genes to ensure a fair comparison among methods. We observed that genes ranked highly by pLI are significantly more likely to be ranked highly by RVIS, and vice versa (~50% mutual overlap for top 3000 genes from each set), consistent with the common origins of both metrics in population-based sequencing data (Fig. 2A). In contrast, fewer genes with high EDS scores are ranked within the top 3000 genes by either pLI or RVIS (~25% vs. 50% overlap between pLI and RVIS, Fig. 2A, **Fig. S4**).

**Figure 2.**
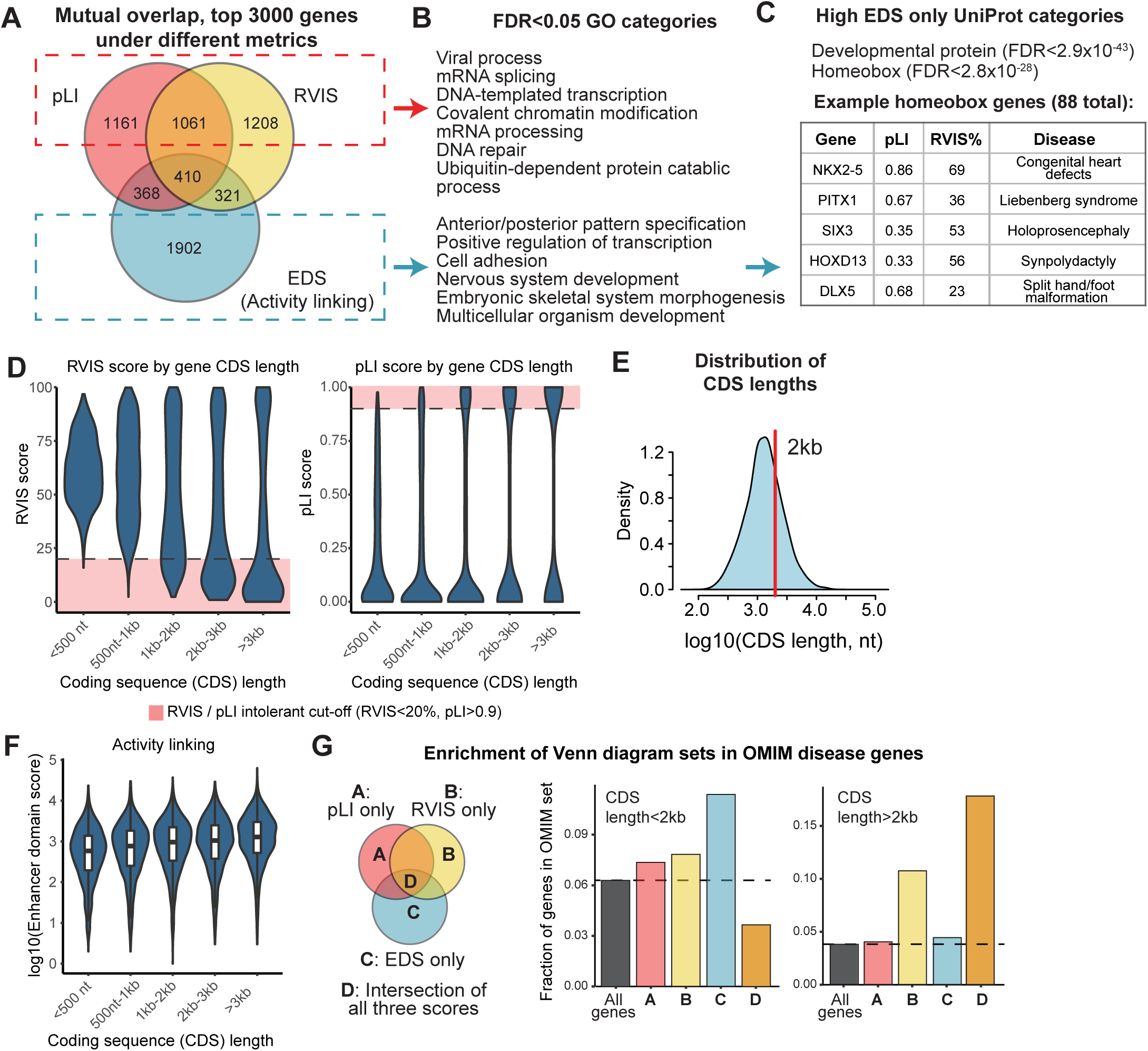
Enhancer domain score is complementary to gene constraint metrics. **(A)** Venn diagram overlap of top 3000 genes ranked by pLI, RVIS and enhancer domain scores. High EDS genes sum to 3001 due to a tie. **(B)** Top GO categories enriched in pLI/RVIS-only gene sets (top, red) and EDS-only gene sets (bottom, blue). Full list of GO enrichments listed in Tables S2 and S3. **(C)** Top, Top enriched UniProt categories in EDS-only gene sets. Bottom, examples of homeobox transcription factors with high EDS rank and low pLI/RVIS ranks (outside top 3000 for both pLI and RVIS). **(D)** RVIS and pLI scores are strongly dependent on gene coding sequence (CDS) length. Shaded area corresponds to commonly applied cut-offs for gene intolerance. **(E)** Distribution of CDS lengths for protein coding genes. Red line corresponds to 2kb, with 65% of human genes less than 2kb in length. **F.** The EDS metric is less correlated with CDS length (x-axis) than pLI/RVIS using activity-linking enhancer domain scores (y-axis). **G.** Left, Venn diagram groups of gene sets corresponding to bars in bar plot. Middle, EDS (activity-linking) in genes with CDS length < 2kb is more predictive than pLI or RVIS for genes in OMIM database. Right, In genes with CDS length > 2kb, greatest enrichment for OMIM genes observed when considering genes ranked highly by all three metrics (pLI, RVIS and EDS).

GO enrichment of genes unique to either pLI/RVIS or EDS reveals that the former gene set (high pLI or high RVIS, low EDS) is enriched for core, housekeeping cellular functions including mRNA processing, DNA replication and cell division (Fig. 2B, **Table S2**). These genes are essential to cellular function, but not involved in a specific human developmental process. In contrast, the latter gene set (low pLI and low RVIS, high EDS) is enriched for genes involved in pattern specification, embryonic development and organ development GO categories (Fig. 2B, **Table S3**). As a particularly striking example of the discrepancy between EDS and constraint-based metrics, we noticed that genes with high EDS but poor pLI/RVIS scores are significantly enriched for the “Homeobox” UniProt category (Fig. 2C, FDR<2.8<10^−28^), including members of the HOX, IRX, NKX and PITX families (**Table S3**). Many of these genes have well-documented roles in congenital disease, including NKX2-5 for congenital heart disease, HOXD13 for synpolydactyly and PITX1 in limb malformations^36–38^ (Fig. 2C). In aggregate, these 88 homeobox genes have a median pLI of 0.43 and RVIS percentile of 47%, dramatically weaker than the commonly-applied cut-offs of 0.90 for pLI and 20% for RVIS, illustrating that these are not borderline genes that were narrowly missed by pLI and RVIS (Fig. 2C).

We investigated the reason why many critical developmental transcription factors and other congenital disease genes would rank poorly by pLI and RVIS, and noticed that gene length is a major factor behind this discrepancy (Fig. 2D). In particular, 65% of all human genes have coding sequences less than 2kb in length (~667 amino acids, Fig. 2E). These short genes have 3.1-fold and 4.4-fold fewer pLI/RVIS-defined highly constrained genes compared to long genes with coding sequences longer than 2kb (p=4.44×10^−272^ for pLI, p<2.2×10^−308^ for RVIS, Fisher’s exact test). This is likely due to the increased statistical power present for long genes to identify a depletion of LoF and missense variants in comparison to expectations. This length-dependent effect is especially pronounced for genes with coding sequences less than 1kb in length, where only 5.0% and 1.3% of genes reach the pLI and RVIS intolerance thresholds, compared to 35.6% and 46.2% for long genes, respectively. In contrast, we observe that while EDS is also associated with gene length (Fig. 2F, **Fig. S4B**), the effect is more attenuated than pLI and RVIS scores.

To further investigate the effect of gene length on disease gene prioritization, we compared the enrichment of OMIM disease genes for different gene sets from Fig. 2A. For shorter genes (coding sequence < 2kb), EDS is the only informative metric for OMIM gene enrichment (group “C” vs. groups A, B, D in *middle* panel of Fig. 2G), while for long genes, the greatest enrichment of OMIM genes is observed for the set of genes that rank highly by all three metrics (group “D” in *right* panel of Fig. 2G). Collectively, these results demonstrate that the EDS is an informative metric for prioritizing disease genes, and that the EDS is complementary to population-based constraint metrics such as pLI and RVIS. Of particular importance to interpreting variation in patient genomes, the EDS metric provides an approach for recognizing disease-causing genes that are too small to provide sufficient guidance about pathogenicity using intolerance and constraint scores based on currently available population genetic data.

### Enhancer domain scores in individual organs reflects specific disease phenotypes

The activity of an enhancer is often restricted to a small set of tissues. We therefore reasoned that the number of functional nucleotides in tissue-specific enhancers could predict the affected tissues for developmental disease genes. In contrast, pLI and RVIS are metrics that reflect the effects of purifying selection at the level of the organism and currently cannot be adapted to generate tissue-specific constraint scores. To calculate a tissue-specific enhancer domain score, we split the tissues used to calculate the original EDS into 18 “tissue groups” that reflect different organs of the body (see **Table S4** for grouping). We counted the number of conserved regulatory nucleotides for tissue-specific enhancers present in each of the 18 tissue groups to obtain a tissue-specific EDS (**Fig. S5A**, see Methods). Any individual tissue group will have fewer linked enhancers than the full set of 127 tissues. Indeed, a median of 2,354 genes in the 18 tissue groups were assigned a tissue-specific EDS, in contrast to 19,038 genes for the original EDS (**Fig. S5B,C**). Given the greater influence of noise, we only considered the top 250 genes (top ~10%) ranked by tissue-specific EDS per tissue group as a proof-of-concept analysis.

To assess whether the tissue-specific enhancer domain score is informative, we considered the affected organs for diseases involving the top 250 genes of each tissue-specific EDS. We relied on annotations from the DDG2P database that includes clinician-curated information on the organ specificity of developmental diseases^31^. We observed that genes involved in diseases affecting different organs are enriched for the corresponding tissue-specific EDS. For example, genes with associated musculature phenotypes are significantly more likely to rank highly by the skeletal muscle-specific EDS. Additionally, genes with associated “heart/cardiovascular/lymphatic” phenotypes are significantly more likely to rank highly by the heart and fat-specific EDS (**Fig. S5D**). These results offer a demonstration that the count of functional nucleotides within tissue-specific enhancers can reflect individual organs affected by disease genes.

### The enhancer domain score reflects resilience against genetic perturbations and complexity of gene expression

We investigated three possible explanations for the strong relationship between EDS and disease-relevance: genes with larger regulatory domains (i) have more complex spatiotemporal gene expression patterns, (ii) are more resistant to environmental perturbations, or (iii) are more resistant to perturbation from genetic variants.

First, we investigated the spatiotemporal expression patterns of high EDS genes by considering the number of tissues or cell types in which they are expressed. We expect that genes with the least complex expression patterns would be ubiquitously expressed (i.e. promoter-driven constitutive expression), while those with more complex expression patterns might be expressed in an intermediate number of tissues, reflecting more precise gene expression regulation. We considered the 18 tissue groups from the Roadmap Epigenomics Project from above, and observed that genes with higher EDS bins are indeed less likely to be ubiquitously expressed, and are more likely to have expression in an intermediate number of tissues (**Fig. S6A,B**). As many of these 18 tissue groups are adult tissues and could behave differently from earlier developmental time points, we also considered two sets of fetal tissues: 1) a subset of 8 embryonic tissues from the Roadmap Epigenomics Project samples, and 2) 87 cell type clusters identified from single-cell sequencing of 50+ mouse organs, tissues and cell lines at different developmental time points^39^. Both developmental gene expression datasets likewise indicate that fewer high EDS genes have ubiquitous expression, and are instead more likely expressed in an intermediate number of tissues (**Fig. S6C-F**). These results suggest that the enhancer domain score is in part connected with a gene’s spatiotemporal gene expression pattern.

Second, we investigated whether genes with greater EDS are more resistant to environmental perturbations. For this analysis, we quantified the stability in gene expression across human individuals. We used data from the GTEx consortium^40^, which has generated RNA-seq data collected from ≥70 adult human individuals in 48 different human tissues (range: 70 to 491 individuals per tissue). Surprisingly, stability in gene expression is inversely correlated with EDS, as genes with the largest regulatory domains have the most variable expression across individuals (**Fig. S7A**). As the GTEx consortium profiled only adult post-mortem samples, we also considered the possibility that expression of high EDS genes is highly constrained at earlier developmental time points. We therefore compiled gene expression data from panels of human iPS-derived cardiomyocytes (42 individuals^41^) and iPS-derived sensory neurons (51 individuals^42^). Both iPS-derived cell types have been reported to exhibit fetal-like gene expression patterns^42,43^. We observed the same trend as seen in adult tissues where high EDS percentile genes have greater expression variability, suggesting this is a general, rather than adult-specific, trend (**Fig. S7B,C**). Together, these data suggest that larger regulatory domains do not promote more stable gene expression patterns, but instead that high EDS genes have greater variability in gene expression across individuals.

Finally, we investigated whether a greater number of functional regulatory nucleotides makes a gene more resistant to expression perturbation from genetic variants. We quantified the proportion of eGenes, which are genes affected by expression quantitative trait loci (eQTLs) that in consequence show allele-dependent differences in expression across human individuals^40^. Under a null model with no regulatory buffering, we would expect that high EDS genes are more likely to be eGenes, owing to their larger regulatory elements that have more opportunities to overlap disruptive genetic variants. Instead, in each of the 48 GTEx tissues, we observed that high EDS genes are ~20% less likely to be eGenes than all genes (p=4.16×10^−24^, paired t-test across 48 tissues, Fig. 3A,B, **Fig. S8A,B**). We observed a similar trend for cell lines resembling earlier developmental time points using eQTL data from iPS-derived cardiomyocytes and iPS-derived sensory neurons (**Fig. S9**), indicating this trend is not specific to adult tissues. We also compared the coefficient of determination for predicting gene expression across GTEx individuals using local, common genetic variation (i.e. elastic net R^2^ in PrediXcan), and observed that high EDS genes have significantly weaker R^2^ values than all genes (Figs. 3D-F, **S10** and **S11**). Together, these results suggest that a high regulatory nucleotide count buffers a gene’s expression against the effects of genetic variation.

To explore the mechanism by which high EDS genes are buffered against regulatory genetic variation, we hypothesized that the enhancer domain score reflects, at least in part, the degree of regulatory redundancy available for a gene. To test this possibility, we quantified the degree of enhancer redundancy per gene by calculating a pairwise Jaccard index for enhancer activity across 127 human tissues between all pairs of enhancers regulating a gene (Figs. 3G and **S12A**). Consistent with our hypothesis, we observe that genes with a high pairwise Jaccard index (i.e. the gene has multiple enhancers with similar activity patterns across tissues) are less likely to be eGenes, across a wide range of EDS bins (Figs. 3H, **S8D** and **S12B,C**). These results suggest that the relationship between EDS and eGenes is due at least in part to the evolution of redundancy of regulatory sequence.

**Figure 3.**
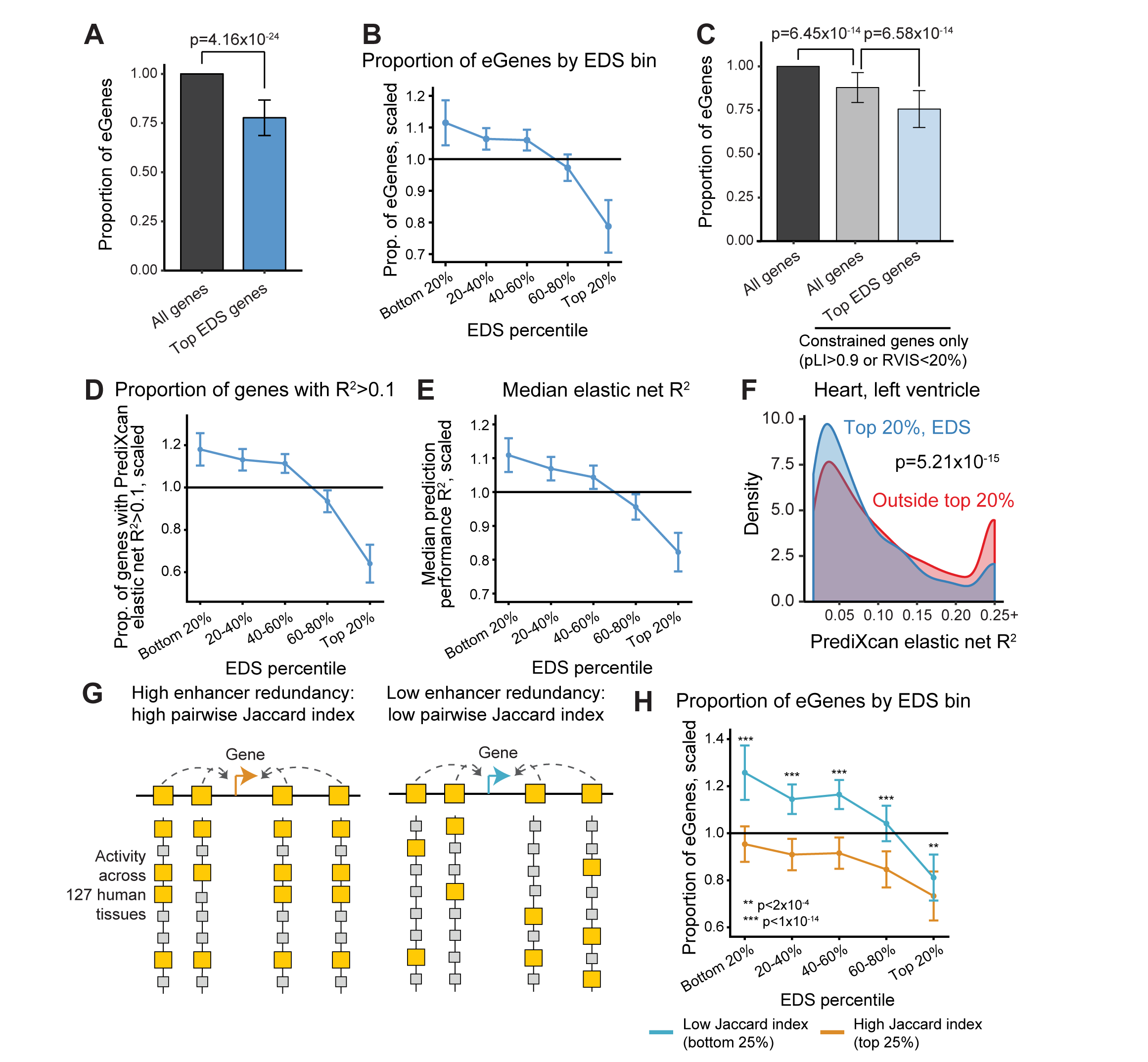
Genes with high EDS and redundant enhancer domains are resilient to genetic perturbations. **(A)** Top 3000 EDS genes are significantly depleted for eQTL eGenes from the GTEx project. Error bars correspond to standard deviation of depletion (normalized to the set of “All genes”) across 48 GTEx tissues with n≥70 samples. **(B)** eGene depletion depends on a gene’s EDS ranking, with the top 20% of genes most depleted. Error bars represent standard deviation across 48 GTEx tissues tested. All values are scaled to the mean rate of eGenes across percentile bins for each tissue, as GTEx sample sizes, and therefore statistical power, differ per tissue. **(C)** Depletion of eGenes in high EDS gene set persists after restricting to highly constrained genes. Error bars correspond to standard deviation across 48 GTEx tissues. p-values from Fisher’s exact test. **(D)** Inverse relationship between EDS percentile and PrediXcan R^2^ for gene expression using GTEx dataset across 48 tissues. Proportion of genes with predicted gene expression R^2^>0.1 by elastic net regression in Wheeler *et al*. 2016 using local common genetic variation. **(E)** Inverse relationship between EDS percentile and median PrediXcan elastic net R^2^ of genes (values from Wheeler *et al*. 2016). Only subset of genes with significant heritability (FDR<0.1) used for calculations. **(F)** Genes in top 20% EDS bin (blue) have lower PrediXcan predicted expression R compared to all other genes (red). Density plots for all 48 GTEx tissues in **Figs. S10** and **S11**. p-value from Mann-Whitney U test for the two full distributions (i.e. without binning genes with R^2^>0.25). **(G)** Identification of genes with high enhancer redundancy by quantification of pairwise enhancer activity patterns across human tissues *Left,* Example gene with high enhancer redundancy. *Right,* Example gene with low enhancer redundancy. **(H)** High enhancer redundancy is associated with reduced rate of eGenes (orange) compared to low redundancy genes (blue). p-values correspond to paired t-test between high and low Jaccard index bins conducted across 48 GTEx tissues, error bars represent standard deviation. Proportion of eGene values were scaled to the mean value of all bins in Fig. 3B

The GTEx consortium previously observed that genes with high LoF intolerance were also depleted as eGenes, and attributed this effect to purifying selection against deleterious regulatory variants^44^. To test these two possibilities (regulatory buffering vs. purifying selection in the regulatory regions of genes), we quantified the depletion of eGenes in the subset of highly constrained genes (pLI>0.9 or RVIS <20%). Consistent with the purifying selection hypothesis, we observed that highly constrained genes have a significantly lower proportion of eGenes compared to all genes (~15% decrease, p=6.45×10^−14^, Fisher’s exact test, Fig. 3C and **S8C**). However, within these highly constrained human genes, genes with high EDS are even less likely to be eGenes than compared to all highly constrained genes, demonstrating a role for regulatory buffering as well (~14% decrease from all highly constrained genes, p=6.58×10^−14^, paired t-test across 48 tissues). These results suggest that regulatory buffering and purifying selection act together to reduce the likelihood of eQTL linkages for high EDS genes.

### eQTLs tend to nominate non-causal genes at GWAS loci

Genetic mapping of complex diseases has revealed that most association signals reside in non-coding DNA regions. As enhancer elements can regulate genes over 1Mb away, the identification of causal genes at these loci remains an outstanding problem. Given that the EDS framework is informative for interpreting rare diseases, we hypothesized that it also applies to complex human traits and diseases. We considered loci from ten genome-wide association studies based on the diversity of affected organ systems and large number of significant genetic loci implicated in each study (>75 loci per study, studies listed in **Table S5**)^45–54^. Consistent with the results we obtained for Mendelian diseases (e.g. Fig. 1E,H), we observed that in nine of the ten GWASs tested, candidate GWAS genes (initially defined as the single nearest gene to each “lead SNP”, commonly selected as the SNP with the strongest p-value) are significantly enriched for high EDS genes, confirming that genes involved in complex human traits and diseases have high enhancer domain scores (Fig. 4A).

As most GWAS loci are non-coding and believed to influence gene expression, many investigators have used eQTL information to prioritize causal genes at individual GWAS loci^45–54^, and have integrated eQTL and GWAS data to perform transcriptome-wide association studies (TWAS) for discovering causal genes genome-wide^55–57^. While applying eQTL information to prioritize biologically relevant causal genes is appealing, our data indicate that (i) GWAS genes have higher-than-normal EDS (Fig. 4A), and (ii) genes with high EDS are depleted from being eGenes (Fig. 3). Furthermore, GWAS loci tend to have more SNPs in linkage disequilibrium^14^, leading to a greater likelihood that the locus is also associated with additional eQTL regulatory signals unrelated to the trait of interest. When considered together, these observations suggest that eQTL prioritization approaches could be prone to implicating wrong genes with low EDS due to regulatory buffering at true causal genes.

**Figure 4.**
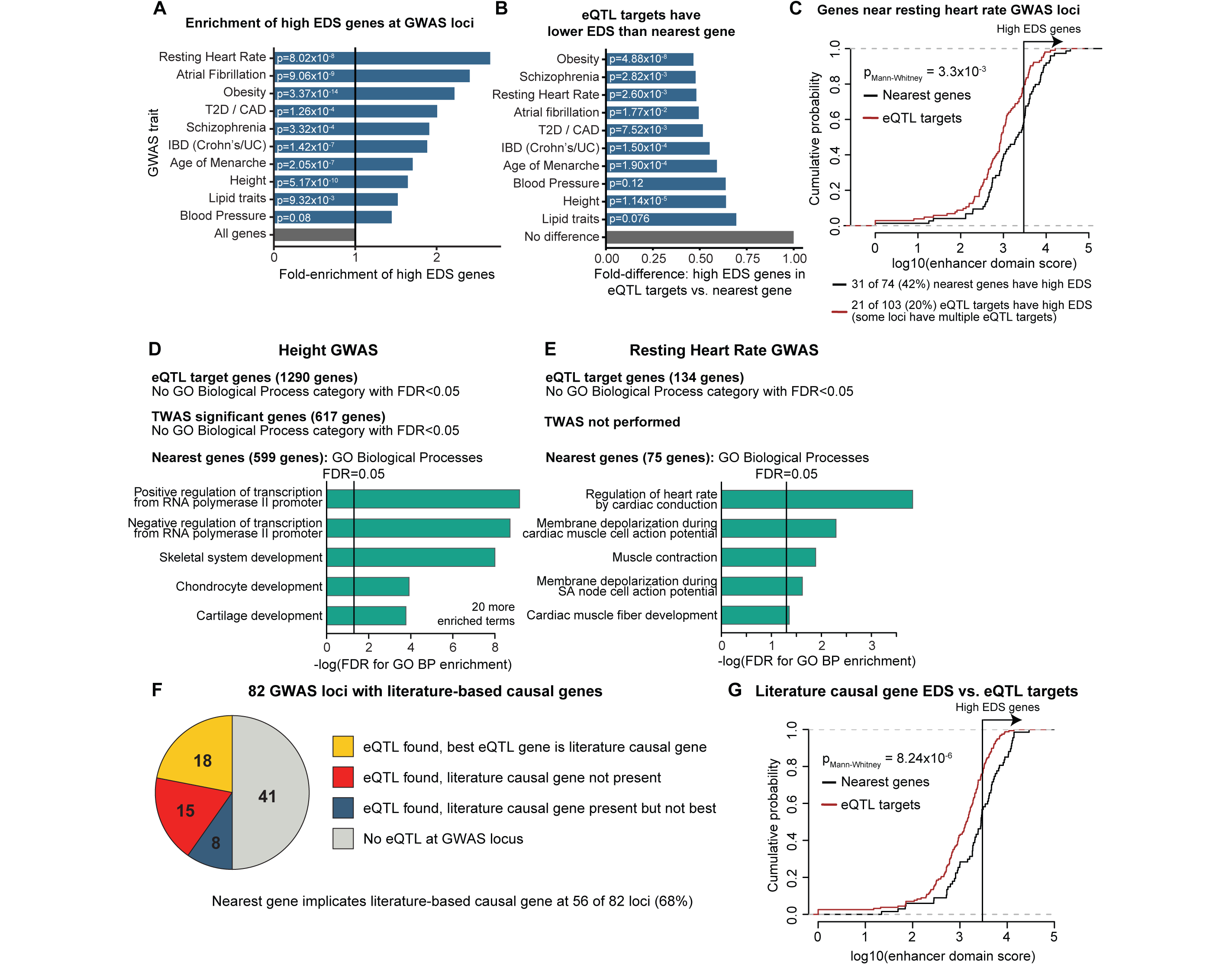
Candidate causal genes at GWAS loci have high EDS and are different from eQTL targets. **(A)** Candidate GWAS genes from ten GWAS studies are enriched for high EDS genes. Candidate GWAS genes were identified by selecting the single nearest gene to the lead SNP at each GWAS locus. p-values are from Fisher’s exact test of the proportion of high EDS genes within candidate GWAS genes compared to all genes. **(B)** eQTL targets at GWAS loci have lower EDS scores than nearest genes. Comparison of proportion of high EDS genes within the set of eQTL target genes at GWAS loci vs. nearest genes. p-values from Fisher’s exact test of proportion of high EDS genes in each gene set. **(C)** Example comparison of the entire distribution of EDS scores for nearest genes vs. eQTL targets for one GWAS study (resting heart rate). p-value from Mann-Whitney U test of EDS scores for genes in each gene set. Pseudocount of 1 added before log transformation. **(D,E)** Significantly enriched GO biological process terms for sets of nearest genes, eQTL target genes and TWAS significant genes for two GWAS studies. Full list of all enriched GO terms for ten GWAS studies considered are available in **Table S5**. **(F)** Literature-based causal genes at GWAS loci are mostly not eQTL targets. eQTLs were identified from the GTEx project and selected from any available tissue. Best eQTL gene was selected on the basis of the strongest p-value. List of loci and correct assignments available in Table S5. **(G)** Literature-based causal GWAS genes have higher EDS scores than eQTL targets of the GWAS loci. p-value from Mann-Whitney U test. Pseudocount of 1 added before log transformation.

To test this possibility, we first compared the EDS scores of eQTL target genes at GWAS loci against the EDS scores of nearest genes, the default but imprecise approach for assigning genes to each GWAS locus^26,28^. Consistent with our hypothesis, the EDS scores of eQTL target genes were significantly lower than the nearest genes (fold difference < 1 for all ten GWASs, p<0.05 for 8/10 GWASs, Mann-Whitney U test). This trend persisted when only considering eQTLs identified in specific disease-relevant tissues (**Fig. S13**). These data indicate that eQTLs have a tendency to implicate low EDS genes (Fig. 4B,C).

The discrepancy between nearest gene targets and eQTL targets can be due to two effects:

i. Current eQTL studies often implicate incorrect, non-causal genes. True causal genes have higher EDS scores and therefore require greater eQTL sample sizes to detect.
ii. Current eQTLs studies tend to implicate the correct causal genes. These causal genes currently identified by eQTLs are biased towards low EDS because these are most easily identified at current levels of statistical power.

We reasoned that if scenario #2 is true, GO enrichments of eQTL target genes will yield more biologically relevant categories than GO enrichments of nearest genes to GWAS loci, as the nearest gene approach is recognized to be imprecise^26,28^. In contrast, under scenario #1 we expect GO enrichment results of eQTL target genes to be less biologically relevant. Across the GWAS studies we tested, we observed a consistent trend where eQTL target genes have weaker or no enrichment for disease-relevant GO categories when compared to nearest genes, consistent with scenario #1 (eQTLs implicate non-causal genes) and suggesting that eQTL target genes are generally not biologically relevant (example GO enrichments in Fig. 4D,E, full list of GO enrichments for all ten GWAS traits in Table S5). We also observed that genes with significant associations in transcriptome-wide association studies (which integrate GWAS and eQTL information) have low EDS distributions and are likewise not enriched for disease-relevant GO categories (Fig. 4D, **Fig. S14, Table S5**)^56^. Together, these results suggest that eQTLs and eQTL-based techniques often nominate non-causal genes at GWAS loci.

To further investigate whether eQTLs implicate the correct causal genes, we compiled a list of 68 literature-based causal genes at 82 GWAS loci selected from all published GWAS studies (**Table S5**). These loci were chosen if the candidate causal gene satisfied one of the following three criteria: (i) the gene is implicated in the Mendelian form of the GWAS trait in OMIM (e.g. chondrodysplasia for height, long QT syndrome for cardiac QT interval length, Wolfram Syndrome for Type II Diabetes), (ii) the gene was implicated at the GWAS locus by a focused experimental study (e.g. IRX3/5 at rs1421085/FT0 obesity locus), or (iii) the gene is targeted by a therapeutic that treats the disease studied in the GWAS (e.g. HMGCR for cholesterol, DRD2 for neuropsychiatric disease). We further excluded loci where the lead SNP or any SNP in strong linkage disequilibrium (r^2^>0.8, CEU cohort from 1000 Genomes project) overlapped a protein-coding exon of the candidate causal gene, as these loci could act through non-regulatory mechanisms.

At these 82 GWAS loci, the literature-based causal gene is the top eQTL target (by p-value) in only 22% of cases, and is one of the targets (FDR<0.05 in GTEx project, any tissue) in 32% of cases (including the 22% where it is the best hit, Fig. 4F). eQTL information is incorrect for 18% of loci where the literature-based causal gene is not an eQTL target, but a different gene at the locus is. In contrast, assigning the nearest gene to each GWAS locus correctly identified the literature-based causal gene at 68% of loci (**Table S5**). Finally, we note that literature-based causal genes have higher EDS scores than eQTL targets of the 82 GWAS loci (Fig. 4G), further indicating that eQTLs preferentially implicate non-causal, low EDS gene targets, rather than the probable correct causal gene. In summary, our results suggest that the genes most relevant for disease have higher EDSs and are consequently less likely to be implicated by eQTLs.

### High burden of rare variants within regulatory regions associates with allele-specific expression

The importance of common non-coding variants in complex human diseases has been well established in recent years^12–14,58^. In contrast, rare variant mapping studies have generally been unable to identify significant disease association signatures in non-coding regions, despite an expectation that many such signals exist^18–21^. A prediction from the EDS analyses presented thus far is that disease genes generally have larger enhancer domains, and will therefore be protected from rare non-coding point mutations through the same regulatory buffering mechanisms presented above. In consequence, we would expect that an effect on gene expression may depend on the presence of multiple point mutations falling in the regulatory regions of a gene.

**Figure 5.**
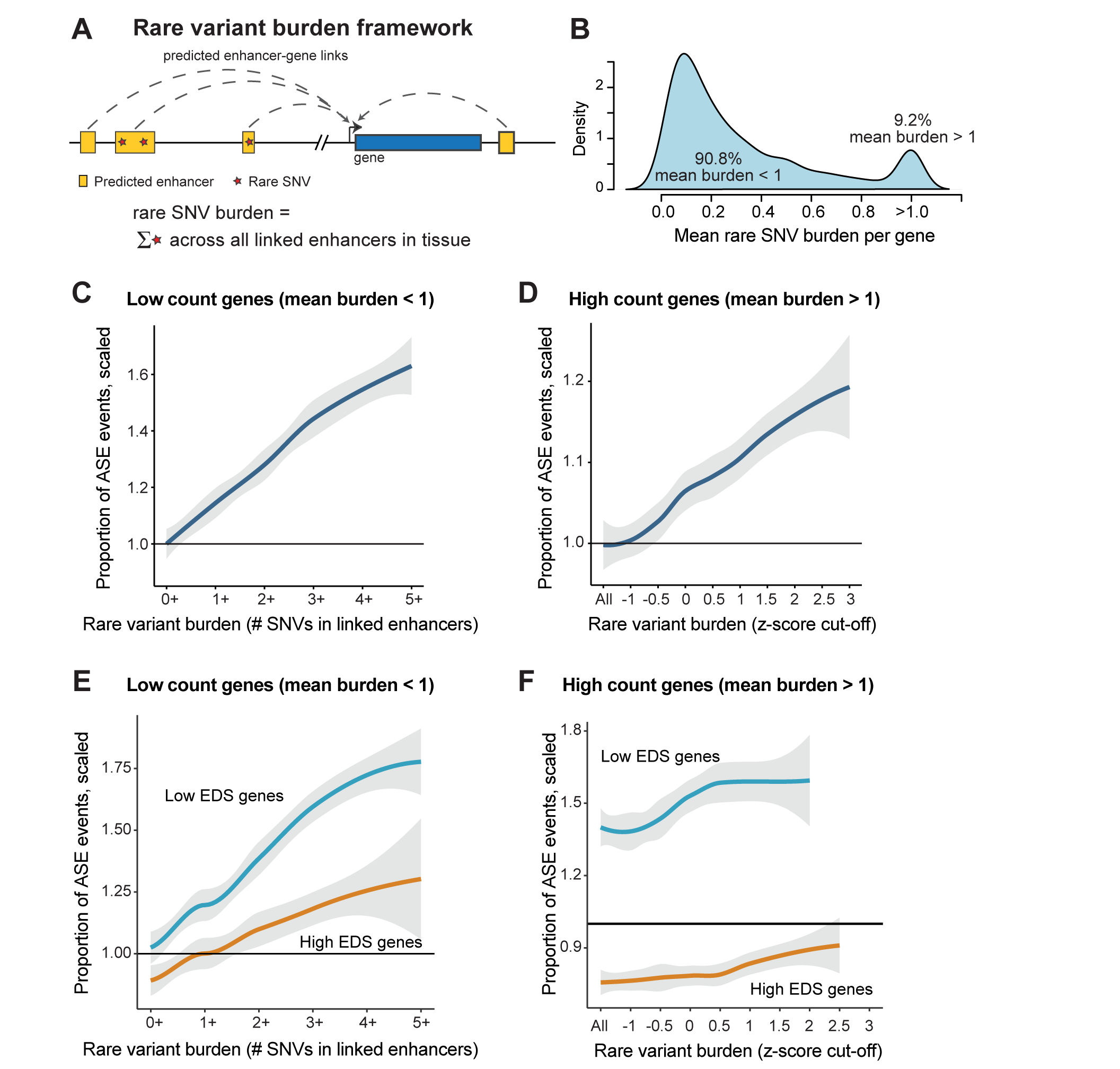
Burden of rare SNVs in regulatory regions is associated with increased rate of allele-specific expression. **(A)** Overview of framework for conducting rare SNV burden analyses. Briefly, the number of rare SNVs (MAF<0.01) within enhancer elements active in a tissue are counted and compared against the allele-specific expression of the gene. All analyses in Fig. 4 performed using activity-linking. Proximity-linking results are presented in Supplementary Fig S11. **(B)** Mean number of rare SNVs per gene across individuals. Analyses in panels C-F are performed separately for genes where mean burden across individuals is less than or greater than 1. **(C,D)** Greater burden of rare SNVs is associated with higher rates of ASE events using enhancers linked to genes by the activity-based linking method. Curve corresponds to Loess curve across GTEx tissues, shaded region represnts 95% CI. **(E,F)** Genes with high EDS scores (orange, top 3000 by EDS) have lower rates of ASE events at same rare variant cut-off numbers than genes with low EDS (all genes outside top 3000, blue). ASE values are scaled to the set of all genes (panels C and D). Curve corresponds to Loess curve across GTEx tissues, shaded region represents 95% CI.

To test this prediction, we first asked whether a high burden of rare non-coding single nucleotide variants (SNVs) in regulatory regions can perturb gene expression, as measured by the presence of allele-specific expression. We hypothesized that while environmental perturbations would often lead to biallelic changes in expression, the genetic perturbation of a *cis*-acting regulatory element should lead to a monoallelic change in gene expression. These monoallelic expression patterns can be identified by detection of allele-specific expression, based on the imbalance of transcript expression in individuals with a heterozygous exonic variant. We developed a framework to compare the burden of rare SNVs (MAF<0.01) in enhancer elements linked to genes against the ASE rate (Fig. 5A,B, **Table S4**). The GTEx project offers an ideal dataset to apply this framework, with 148 individuals profiled by both whole-genome sequencing and RNA-seq across 48 tissues (GTEx v7). We observeda that an increased SNV burden is associated with greater ASE rates in a dose-dependent manner (Fig. 5C,D, **Fig. S15A-C**, see Methods for details on analysis). Notably, this relationship is strongest when considering the SNV burden across the entire predicted enhancer elements (both conserved and nonconserved nucleotides), and is attenuated when considering only the SNV burden in conserved nucleotides (**Fig. S16**), indicating that genetic variants at non-conserved regulatory nucleotides can disrupt gene expression. In subsequent analyses, we focus on the SNV burden in the entire enhancer elements.

As a control, we shifted the position of the linked enhancers used in our analyses by 200kb and 500kb both upstream and downstream from their original positions, to maintain the total number of regulatory nucleotides linked to each gene. In all cases (different shifting windows, gene sets tested, and linking methods), the SNV burden in the shifted enhancers was more weakly associated with the ASE rate than the true set of linked enhancers (**Fig. S17**). Together, these results indicate that a high SNV burden in enhancers is more likely to result in allele-specific expression of the linked gene, and suggests that quantifying regulatory burden in disease studies can aid in identification of potential causal genes.

Finally, we tested whether the expression of high EDS genes is less likely to be disrupted by a high SNV burden. We split genes into high and low EDS groups (top 3000 EDS genes vs. all genes outside the top 3000), and observed that the burden-ASE curve is attenuated for high EDS genes, indicating that nearby high EDS genes, a greater SNV burden is needed to achieve the same rate of transcriptional disruption as low EDS genes (Fig. 5E,F, **Fig. S15D,E**). In summary, these results indicate that high EDS genes, which are enriched in human disease, are more resistant to regulatory perturbation.

## Discussion

In this study, we show that the number of functional nucleotides predicted to regulate a gene is closely related to the gene’s importance in development and disease. This nucleotide count, which we call the “enhancer domain score”, is predictive for disease-causing genes and provides information independent of existing commonly-used metrics of gene essentiality, including pLI and RVIS. When EDS differs with these population-scale metrics of gene constraint, EDS is often more effective at identifying developmental and disease genes, especially in genes with short coding sequences. We show that genes with a high EDS, and especially those with redundant enhancer elements, are depleted for eQTLs and have evolved robustness to regulatory variation. To illustrate the implications of this relationship, we focus on the problem of identifying causal disease genes from GWAS studies. We show that candidate causal genes at GWAS loci have high EDS and are distinct from eQTL targets of the same loci, suggesting that using eQTL overlaps to prioritize causal GWAS genes can be misleading. Furthermore, by showing a significant relationship between EDS and disease-causing genes and between the burden of variation in linked enhancers and gene expression, we establish a framework to interpret the burden of rare non-coding signals from whole-genome sequencing studies. Collectively, these results provide new insights into the identification of disease genes, as well as the disruption of gene regulation by regulatory variants.

Recent studies indicate that the majority of genetic variants that disrupt TF binding or chromatin modifications at enhancer elements do not have an effect on gene expression^9–11^. This observation has been attributed to a number of factors, including “futile” regulatory activity and regulatory redundancy^11^. Our results support the role of regulatory redundancy to explain this discrepancy, and in particular highlights that disease-relevant genes have large enhancer domains and will be most buffered against genetic variation. A number of recent studies have experimentally mapped enhancer-gene regulatory links by using CRISPRi to experimentally inactivate enhancers and profiling gene expression differences^25,59^. A prediction from our results is that the expression of disease-relevant genes will be buffered against strong expression effects due to the inactivation of individual enhancer elements, and that the detection of enhancer-gene links for disease-relevant genes will require substantially higher sensitivity or multi-enhancer inactivation approaches. Indeed, in a recent preprint, Gasperini *et al.* found that housekeeping and non-housekeeping genes are disrupted at equal rates by CRISPRi-based enhancer inactivation^59^, seemingly in contradiction to results showing that non-housekeeping genes are regulated by more enhancer elements and should therefore be more likely to be affected^4,5^, which the authors hypothesized could be due to the effects of redundant shadow enhancers.

The computational approaches we used to link enhancers to genes are perceived to have substantial false positive and false negative rates^26–28^. Thus, it is striking that the enhancer domain score we calculated is effective for predicting disease genes. One explanation for the success of the EDS is that while any individual predicted enhancer-gene link is unreliable, aggregation of all predicted enhancer-gene links across 127 human tissues results in a robust and informative score. We also observed that enrichment for disease genes is highest when we count the number of evolutionarily conserved nucleotides, but occurs when considering other metrics that reflect enhancer functionality, including the number of all enhancer nucleotides, the number of nucleotides in TF binding sites, and number of discrete enhancer elements that regulate a gene. We believe that the success of the EDS using conserved nucleotides therefore reflects filtering for high-confidence enhancer elements, as the majority of computationally-predicted enhancer elements do not display activity *in vivo^60^*. As high-quality genome-wide predictions of enhancers and enhancer-gene links become available, the accuracy of the enhancer domain score and tissue-specific EDS can be improved, and non-conserved nucleotides can also be incorporated into the enhancer domain score. Other regulatory elements, including promoters and open chromatin sites can also be included alongside enhancers to construct a comprehensive regulatory score. Additionally, a machine learning model can be trained to incorporate additional regulatory features, including the number of enhancers, pairwise Jaccard index, and the site-frequency spectrum of genetic variation at regulatory regions.

Our observation that candidate GWAS genes have high EDS has implications for using eQTL data to prioritize causal genes. Consistent with our results, other recent studies have also noted the limitations of using eQTL information in GWAS analysis^61,62^. While we expect the causal genes at GWAS loci will be eQTL targets with sufficient study sample sizes, our results support a conjecture raised by Hormozdiari *et al.* where the causal eQTL signals at GWAS loci are “secondary signals in comparison to the stronger associations found in current eQTL studies”^63^. As eQTL study sample sizes grow and the causal GWAS eQTL interactions are detected, additional non-causal signals will concurrently be discovered that obscure prioritization of the causal gene. Co-localization analyses^61,63,64^ that aim to identify instances where GWAS and eQTL loci share the same causal genetic variant should therefore be critical for prioritizing causal genes using eQTL data. Recent co-localization studies have noted a limited causal variant overlap between GWAS and eQTL signals, supporting our data suggesting that most eQTL signals at GWAS loci (identified without co-localization of causal SNPs) are non-causal. However, the co-localization approach may be more fruitful when eQTL studies have larger sample sizes and are performed under additional environmental conditions and with more cell types.

Finally, our observation that high EDS genes are more resilient against regulatory perturbations has implications for the discovery of causal non-coding rare variants in human disease studies. Recent whole-genome sequencing studies have had limited success in identifying rare noncoding single nucleotide variants associated with disease, in particular when focusing on individual regulatory elements^18,19^. The prevalence of regulatory buffering in disease-relevant genes suggests that a burden framework will be most appropriate for discovering significant signals in the non-coding genome, and suggests that enhancer-gene linking approaches are already suitable for identifying the regions within which to develop gene-specific regulatory burden scores. Our results also help explain the longstanding puzzle of why dosage-sensitive genes that cause rare human diseases have many pathogenic mutations in exons but so few point mutations in regulatory regions. Collectively, our work implies that the focus of attention for regulatory variation should encompass a burden approach for point mutations and implies that other mutational classes may be of particular importance, such as short tandem repeats and structural variants.

## Acknowledgements

We wish to thank Abhishek Sarkar and Hae Kyung Im for very helpful discussions and Athma Pai, Sarah A. Dugger, Michael Wainberg, Nasa Sinnott-Armstrong, and Andrew S. Allen for helpful comments on the manuscript. This work was supported by funds from NIH U01 MH105670 and a genome sequencing and analysis grant from Biogen.

## Author Contributions

X.W. and D.B.G. conceived of the study and wrote the manuscript. X.W. performed analyses, D.B.G. secured funding.

## Methods

### Calculation of the enhancer domain score

To maximize the set of transcriptional regulatory elements across tissues, we considered predicted enhancer elements from the Roadmap Epigenomics Consortium^65^. Enhancer predictions from the 15-state ChromHMM model are available for 127 human tissues/cell types and were downloaded from egg2.wustl.edu/roadmap/web_portal. We used two computational approaches to predict enhancer-gene interactions: activity-linking (primary method used in main figures) and proximity-linking (independent approach that treats each tissue individually and does not use RNA-seq information, used in Supplementary Figures). Activity-linking-based enhancer-gene links using the 15-state ChromHMM model across the same 127 human tissues were obtained from Liu *et al.* 2017^29^ (code originally developed in Ernst *et al.* 2011^4^) and were downloaded from www.biolchem.ucla.edu/labs/ernst/roadmaplinking (“RoadmapLinks.zip” file). Proximity-based enhancer-gene links were generated using GREAT v3.0.0 webtool (great.stanford.edu/public/html/index.php^33^), using default settings (“Basel plus extension” linking method, 5kb upstream, 1 kb downstream, plus distal up to 1000kb). We ran GREAT after splitting the set of ChromHMM Epigenome Roadmap enhancers into bins of 500,000 enhancers each.

To calculate an enhancer domain score across all tissues, we used mergeBed (BEDTools v2.26.0^66^) to merge the set of regulatory elements linked to each gene. We considered evolutionarily conserved nucleotides identified by phastCons for three comparisons: phastCons100way, phastCons46wayPlacental and phastCons46wayPrimates. To count the number of evolutionarily conserved nucleotides, we downloaded BED files of evolutionarily conserved elements from the UCSC Genome Table Browser and assigned them to linked enhancer elements using the intersectBed tool (BEDTools v2.26.0). Correlations between enhancer nucleotides and conserved nucleotides (**Fig. S1** and **S3D,E**) were calculated in R, and plots were generated using the hexbinplot function (hexbin package) with the xbins=100 setting. We obtained BED files for cis-regulatory modules and TF binding sites across human tissues from the UniBind database (unibind.uio.no)^30^.

### Enrichment of disease genes in high EDS genes

Sources of disease gene lists used to generate Fig. 1E are listed in **Table S6**. All gene lists were first converted to Ensembl Gene IDs using Ensembl Gene IDs and gene symbols listed in the Ensembl GRCh38 GTF file. The list of organs affected by developmental diseases (used for Fig. 1H) was generated by using grep on the “organ specificity list” column in the DDG2P spreadsheet. We calculated enrichment by considering the proportion of high EDS genes in the gene set, compared to the set of high EDS genes in the entire human genome.

### Comparison of EDS to pLI and RVIS

pLI scores were downloaded from ExAC v0.3.1 (ftp.broadinstitute.org/pub/ExAC_release/release0.3.1/functional_gene_constraint/fordist_cleaned_exac_r03_march16_z_pli_rec_null_data.txt), and RVIS scores were downloaded from RVIS v3 (http://genic-intolerance.org/data/GenicIntolerance_v3_12Mar16.txt), using scores constructed on the ExAC data release. GO enrichments were performed using DAVID v6.8 (https://david.ncifcrf.gov/) with a background set of 20,047 genes that have EDS scores, pLI scores and RVIS scores annotated. All gene lists used for GO enrichment calculations, and resulting lists of enriched GO categories are available in **Tables S2** and **S3**.

### Tissue-specific enhancer domain scores

We grouped 121 of 127 human tissues and cell types from the Epigenome Roadmap project into 18 tissue groups based on groupings provided by the Roadmap consortium (see Metadata at egg2.wustl.edu/roadmap/web_portal/meta.html). Six additional tissue groups (Adrenal, Bone, Cervix, Kidney, Ovary, Spleen) were excluded as only one Roadmap tissue sample mapped per group, which could lead to problems with lack of enhancer sample size and high noise. A list of the groupings, including excluded samples, is available in. To identify tissue-specific enhancer elements, for each tissue group, we merged all enhancers and quantified the number of additional tissues (outside the original tissue group) where the enhancer was present, selecting enhancers active in fewer than 10 additional samples. Tissue-specific EDS scores were calculated following the same process as the organism-level EDS. p-values for enrichment of disease genes within the top 250 genes for any individual tissue-specific EDS score were calculated using Fisher’s exact test comparing to the set of all genes. Multiple testing correction was performed with the Benjamini-Hochberg correction.

### Spatiotemporal gene expression patterns by EDS bins

We used two sets of RNA-seq data across human and mouse tissues to assess spatiotemporal expression patterns for genes: gene expression across 57 human tissues profiled by the Roadmap Epigenomics project (downloaded from the Roadmap Epigenomics portal, https://egg2.wustl.edu/roadmap/data/byDataType/rna/expression/), and mouse single-cell RNA-seq from different time points (embryonic, fetal & adult, downloaded as “MCA_Figure2-batch-removed.txt.tar.gz” from Han *et al*. 2018, available at figshare.com/s/865e694ad06d5857db4b). As the Roadmap Epigenomics dataset samples different organs in different levels of detail (e.g. 10 regions of the adult brain vs. 1 sample for each of kidney, spleen, liver), we grouped tissues into 17 “tissue groups” to avoid biasing our analyses (note, for tissue-specific EDS scores, we used 18 tissue groups. The discrepancy is due to incomplete RNA-seq profiling for Roadmap tissues). As many of the tissue groups represent adult time points, we also considered a separate set of 8 embryonic tissue groups. Tissue-to-group assignments are listed in Table S4. The mouse single-cell RNA-seq dataset we obtained was processed to assign cells into 87 cell clusters from a variety of embryonic, fetal and adult tissues. We generated a single RNA-seq expression vector per cell cluster by summing across all constituent cells, converted mouse gene expression data to human using Ensembl’s BioMart database (mouse gene GRCm38.p6), and for genes with multiple human orthologs, selected the ortholog with highest gene expression. We then converted counts to RPKM and set a minimum cut-off of 10 RPKM.

### Gene expression variability by EDS bins

To measure variability of gene expression, we used three datasets of gene expression across individuals: the GTEx v7 dataset (48 tissues with >80 individuals each), gene expression across iPS-derived cardiomyocytes (Knowles *et al.*, 2017, 42 individuals in control untreated group), iPS-derived sensory neurons (Schwartzentruber *et al.*, 2017, 51 individuals). For GTEx samples, we calculated the coefficient of variation per gene (when expressed, TPM>=1 cutoff) across individuals, and aggregated across tissues using the mean and minimum. Both mean and minimum yielded the same trend (top 20% EDS bin has greatest coefficient of variation), and we show violin plots for mean coefficient of variation in **Fig. S7**. We set minimum expression cut-offs of 10 RPKM for the Knowles *et al.* and Schwartzentruber *et al.* datasets (TPM data was not available), also finding the same trend observed in GTEx samples.

### Proportion of eGenes by EDS bins

We performed eQTL analyses using processed data from 48 tissues generated by the GTEx consortium (v7). Data was downloaded from the public GTEx portal (filename containing eQTL links for all tissues: GTEx_Analysis_v7_eQTL.tar.gz), and eGene lists and significant SNP-gene associations were taken from the *.v7.signif_variant_gene_pairs.txt.gz files. The sample sizes for the 48 tissues ranges from 80 to 491. For each tissue, we considered the number of eGenes discovered within each EDS bin compared to the total number of genes in the EDS bin. Top genes by EDS in Figs. 3A,C and **S8A,C** were chosen as those ranking in the top 3000 by activity and proximity-linking, respectively.

To calculate Pairwise Jaccard indices, we merged all linked enhancers across tissues per gene. For each linked enhancer, we constructed a binary activity vector of length 127, indicating whether the enhancer is active in each of 127 human tissues profiled by the Epigenome Roadmap project. Activity for linked enhancers was defined as overlapping a ChromHMM-called enhancer element (15-state model) in the corresponding tissue. For each gene with more than 3 linked enhancers, we calculated the average pairwise Jaccard index across enhancers using the binary activity vectors per enhancer.

### PrediXcan prediction performance in GTEx by EDS bins

To supplement the eQTL/eGene analyses, we also assessed whether the coefficient of determination R^2^ of the fitted elastic net regression of gene expression on cis-genotypes in GTEx individuals was dependent on a gene’s EDS bin. We obtained R^2^ values from PrediXcan trained on GTEx v7 (available at predictdb.org, under the “download-by-tissue” folder) and used the “pred.perf.R2” variable in the gtex_v7_[tissue]_imputed_europeans_tw_0.5_signif.db files accessed using the RSQLite package in R.

### GWAS analyses

We identified ten GWAS studies with >75 loci from the NHGRI-EBI GWAS catalog^67^ covering traits with a range of ages of onset and affected tissues. In the first part of our analysis, we defined GWAS candidate genes as the single nearest gene to the lead SNP at each locus using the GREAT tool (v3.0.0)^33^. We identified eQTL target genes of these same loci using eQTL data from the GTEx project (v7), selecting significant eGenes using the *_v7.signif_variant_gene_pairs.txt files. We identified eGenes using eQTLs present in any tissue (main analyses), as well as only those in a trait-relevant tissue (used in **Fig. S13**). Significant TWAS genes for each trait were downloaded from Mancuso *et al.* 2017^56^. GO enrichments for all gene sets were performed using DAVID v6.8 (https://david.ncifcrf.gov/). To compile a list of candidate causal genes at GWAS loci, we scanned the literature for instances where loci were nearby a corresponding Mendelian disease gene, where the locus was studied in a focused experimental study, or when the locus is nearby a gene targeted by a therapeutic for the same disease. For each lead SNP in these loci, we identified all SNPs in strong LD (r^2^>0.8, EUR cohort from 1000 Genomes project) and removed the subset of loci that overlapped a protein-coding exon in the putative causal gene (note that loci overlapping other non-causal genes were not removed). After filtering, we were left with 68 literature-based causal genes at 82 GWAS loci.

### SNV burden in regulatory regions

We compiled lists of enhancers active per tissue using the tissue groupings previously used for **Fig. S5** (groupings listed in **Table S4**). For SNV burden analyses, we considered SNVs within the entire enhancer elements, as well as those within evolutionarily conserved regions (merger of phastCons elements across Primates, Vertebrates and Placental Mammals). We preprocessed genotype information from the GTEx project (dbGAP accession phs000424.v7.p2) to select for rare variants. As the GTEx sample size is modest (148 individuals), we used allele frequency information from the BRAVO database (TOPMed Freeze 5, https://bravo.sph.umich.edu/freeze5/hg38/) after mapping BRAVO variants from hg38 to hg19 using the LiftoverVCF tool in Picard tools (v2.9.0). We mapped MAF<0.01 variants to enhancer elements grouped by tissue group, and for each GTEx individual, we counted the number of rare variants per gene, per tissue group for each of the four enhancer sets (activity-linked entire enhancers, proximity-linked entire enhancers, activity-linked conserved enhancer elements, and proximity-linked conserved enhancer elements).

Next, we linked the SNV burden to allele-specific expression of genes. Allele-specific expression information per gene, per tissue and per individual was obtained from the GTEx project (phs000424.v7.p2). For each GTEx individual, we filtered the processed ASE data to create (i) a list of genes per tissue with significant allelic expression (adjusted p-value < 0.05), and (ii) a background list of all genes per tissue tested for allelic expression (i.e. containing a heterozygous SNP). As we wanted to assess whether a high burden of SNVs in linked enhancers was associated with allelic expression, we matched enhancer tissue groups to corresponding tissues profiled by the GTEx project (see **Table S4** for list). In total, we matched 37 GTEx tissues to 14 different enhancer groupings from the Epigenome Roadmap project.

We first noticed that a small number of GTEx individuals had significant allelic expression for a very large number of genes (e.g. one individual had allele-specific expression for 9,462 genes in skin, but ASE for <1000 genes in all other tissues). We hypothesized these could be due to sequencing or data processing artifacts. For each GTEx tissue, we therefore removed the 10 individuals with the greatest number of genes showing significant allelic expression. A small number of genes also consistently showed allele-specific expression across a large number of individuals. A literature search revealed that many of these are imprinted genes (e.g. MEG3, PLAGL1, L3MBTL1) or HLA genes. To avoid potentially confounding signals, we therefore removed genes listed in the Imprinted Gene Catalog (http://igc.otago.ac.nz) and genes identified in a global survey of imprinting (Baran *et al*. 2015^68^). The full list of 109 genes excluded from ASE analysis is listed in Table **S7**. Finally, we also removed samples marked as outliers by the GTEx project (https://www.gtexportal.org/home/documentationPage#staticTextAnalysisMethods).

In considering the distribution of regulatory SNV burdens per gene (e.g. Figs. 5B and **S15A**), we realized that genes with high and low average SNV burdens would need to be analyzed separately. In genes with low average SNV burden, most individuals have 0 or 1 rare SNV in a linked enhancer. In these genes, the raw number of rare SNVs directly reflects the relative burden (e.g. observing 5 SNVs is high if most other individuals have <1). In contrast, the mean SNV burden for high count genes has a wide spread, and SNV burdens need to be z-score transformed before analysis (e.g. observing 5 SNVs in an individual could be either high or low, depending on whether the mean SNV count across individuals is 1.5 or 20). After we split genes into high and low count groups (mean SNV count >1 and <1, respectively), in each tissue we calculated the proportion of gene-by-individual pairings with allele-specific expression at each SNV burden cut-off. As the proportion of genes showing ASE per tissue is different, we merged tissues together to generate the plots shown in Figs. 5C-F and **S15B-E** after scaling each tissue to the ASE rate in all genes (“0+” bin for low count genes and “All” for high count genes). Loess curves and 95% CI were generated using the geom_smooth function in ggplot2.

## Supplementary Tables

**Table S1:** List of genes with enhancer domain scores, pLI and RVIS scores.

**Table S2:** List of enriched GO categories for genes with high pLI/RVIS scores and low EDS

**Table S3:** List of enriched GO categories for genes with low pLI/RVIS scores and high EDS

**Table S4:** List of tissue groupings and matched groupings between Epigenome Roadmap enhancers and GTEx tissues

**Table S5:** GWAS GO enrichment categories and assignment of literature-based causal genes

**Table S6:** Sources of disease gene lists used in enrichment analyses

**Table S7:** List of literature-based imprinted genes removed during ASE analysis

